# Recombinant Pichinde reporter virus as a safe and suitable surrogate for high-throughput antiviral screening against highly pathogenic arenaviruses

**DOI:** 10.1101/2024.12.19.629433

**Authors:** Carolina Q. Sacramento, Ryan Bott, Qinfeng Huang, Brett Eaton, Elena Postnikova, Ahmad J. Sabir, Malaika D. Argade, Kiira Ratia, Manu Anantpadma, Paul R Carlier, Hinh Ly, Yuying Liang, Lijun Rong

## Abstract

Several arenaviruses, such as the Old World (OW) Lassa virus (LASV) and the New World (NW) Junin virus (JUNV), can cause severe and lethal viral hemorrhagic fevers in humans. Currently, no vaccines or specific antiviral therapies are FDA-approved for treating arenavirus infections. One major challenge for the development of new therapeutic candidates against these highly pathogenic viruses is that they are BSL-3/4 pathogens that need to be handled in high biocontainment laboratories. In this work, a recombinant non-pathogenic New World arenavirus, Pichinde virus (rPICV), was used for the development of a high-throughput screening (HTS) assay in the BSL-2 laboratory for the screening and identification of small molecule inhibitors against arenaviruses. The rPICV is a replication-competent virus expressing the firefly luciferase reporter gene in the infected cells proportionally to the infection rate. rPICV infection was optimized for an automated HTS in 384-well format with robust Z′ scores, high signal-to-background ratios, and low intrinsic variance. Screening an established library allowed for the identification of five top hit compounds, which included ribavirin, a known inhibitor of arenaviral RNA synthesis, showing good potency and selectivity in inhibiting rPICV replication. The antiviral activity of the top hit compounds was further validated against another recombinant arenavirus, the OW lymphocytic choriomeningitis virus (rLCMV) and against laboratory strains of LASV (Josiah) and JUNV (Romero). The use of rPICV in the HTS-based antiviral assay under BSL-2 condition has proven to be safe and suitable for the identification of broad-spectrum small molecule inhibitors against highly pathogenic arenaviruses.

## 1. INTRODUCTION

Arenaviruses are rodent-borne enveloped, single-stranded, RNA viruses that can cause a spectrum of diseases including severe and lethal viral hemorrhagic fevers in humans. Lassa virus (LASV) is an Old World (OW) arenavirus found in West Africa and has been estimated to cause 100,000 to 300,000 infections and ∼5,000 deaths annually, according to the US Center for Disease Control and Prevention. The overall estimated fatality rate of Lassa fever is ∼1% but it can be considerably higher in hospitalized patients (∼18%) and during outbreaks (up to 25%) (Roberts, 2024). Apart from causing severe multisystemic disease, LASV infection can also result in severe sequelae such as spontaneous abortion in infected pregnant women (Bello et al., 2016; Khan et al., 2008) and sensorineural hearing loss in one-third of recovered patients (Cummins et al., 1990). In South America, Junin virus (JUNV) and other New World (NW) arenaviruses (e.g. Machupo, Guanarito, Chapare, and Sabia viruses) can cause sporadic outbreaks with fatality rates of up to 30%. Lymphocytic choriomeningitis virus (LCMV), an OW arenavirus with a global distribution, is a neglected human pathogen causing neurological disease, congenital infection, and transplant-associated VHF with substantial mortality.

There are no FDA-approved vaccines or specific treatments against arenaviruses. Although not licensed for treating Lassa fever, ribavirin, a broad-spectrum nucleoside analogue, has been long used to treat Lassa infections (McCormick et al., 1986). However, the clinical evidence gathered through the years is leading to a reevaluation of its clinical benefits (Salam et al., 2022, 2021). New therapeutics for Lassa fever are in various developmental stages. Favipiravir, a purine analog licensed for treating influenza in Japan, showed good efficacy in animal models of severe LASV infection (Lingas et al., 2021; Oestereich et al., 2016), but its efficacy and safety in humans need further evaluation (Raabe et al., 2017). Other small molecule inhibitors targeting viral entry (Iyer et al., 2024) and broadly neutralizing monoclonal antibodies (Li et al., 2022) are in the pipeline and may prove valuable in future clinical trials. Still, Lassa fever is listed as a priority disease on the World Health Organization’s R&D blueprint of vaccines and therapeutics given its severe disease outcomes and pandemic potential (Salami et al., 2020).

Hemorrhagic fever-causing arenaviruses are classified as biosafety level 3 or 4 (BSL-3/4) pathogens and, as such, their handling requires high biocontainment laboratories, imposing major technical challenges for the development of new therapeutic candidates. Prototypic arenaviruses, such as LCMV and Pichinde virus (PICV), have been successfully used as experimental tools in BSL-2 for studying arenaviruses in vitro and in vivo (Amaya et al., 2021a; Lan et al., 2009). PICV is a non-pathogenic NW arenavirus that does not cause disease in humans, but had been adapted to cause Lassa fever-like disease in guinea pigs (Lan et al., 2020, 2009; Shieh et al., 2020). Similar to the other arenaviruses, its genome is composed of two ambisense RNA segments: the small (S) and the large (L) segments, each encoding two genes in opposite orientations (Garry, 2023; Murphy et al., 2024). A stable recombinant PICV with a tri-segmented genome (rPICV) was developed and has been used to study replication-competent Pichinde viral vector-based vaccines against different pathogens, such as influenza and *Mycobacterium tuberculosis* (Dhanwani et al., 2016a; Kirk et al., 2023).

In this work, a high-throughput screening (HTS) assay based on the tri-segmented rPICV reporter virus expressing firefly luciferase (LUC) was developed to identify broad-spectrum antivirals against arenaviruses. Five top hit compounds, which included ribavirin, were identified from the HTS of a library of approved small molecules by measuring LUC enzymatic activity in rPICV-infected cells. These compounds inhibited rPICV and rLCMV with good selectivity, and their antiviral activities were also validated against laboratory strains of LASV and JUNV. Counter-screenings against other viruses besides arenaviruses and time-of-addition assays suggest that the hits may have host- and/or virus-directed antiviral activities by targeting different steps of the rPICV life cycle. These data provide proof-of-concept for the effective use of rPICV as a safe and convenient surrogate system for HTS antiviral assay to identify broad-spectrum small molecule inhibitors targeting various stages of arenaviral life cycle in a BSL-2 environment.

## 2. MATERIALS AND METHODS

The detailed methods are described in the Supplementary Materials and Methods. The most significant aspects are briefly described in this section.

### 2.1. Viruses

The tri-segmented recombinant PICV (rPICV) expressing LUC and green fluorescent protein (GFP) (Fig. 1A) was produced in BSRT7-5 cells, propagated in BHK-21 and titrated in Vero cells by plaque forming unit (PFU) assays as previously described (Dhanwani et al., 2017, 2016b). rLCMV expressing LUC and GFP was created in a similar approach. Working stocks of laboratory strains of LASV (Josiah) and JUNV (Romero) were generated and titrated in Vero cells.

**Fig. 1.**
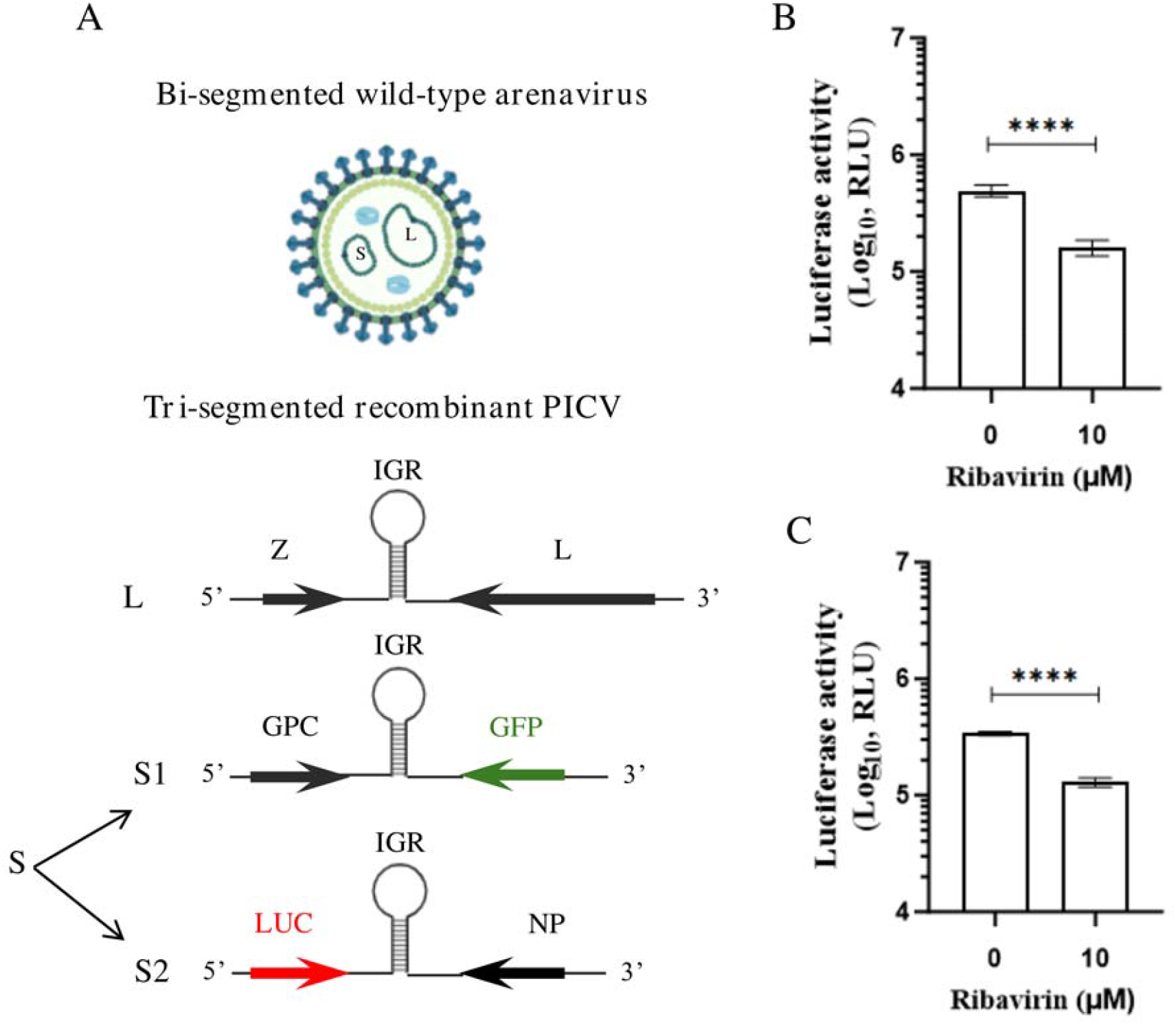
Tri-segmented luciferase-expressing recombinant PICV (rPICV) as tool for antiviral testing. (A) Schematic representation of a wild-type arenavirus with bi-segmented genome (L and S RNA segments) (top) and tri-segmented rPICV genome (L, S1 and S2 RNA segments) encoding the green fluorescent protein (GPF) and firefly luciferase (LUC) as reporter genes (bottom). Luciferase expression was quantified in A549 cells in 96-(B) or 384-well plates (C) infected with rPICV (MOIs of 0.01 and 0.05, respectively) for 24 h. Ribavirin at 10 µM was used as a positive control for inhibition of rPICV replication in both assay formats. Cells were lysed and luminescence was measured using the Neolite Reporter Gene Assay System. Data are presented as the mean ± standard deviations of the relative luminescence units (RLU) measurements from at least three replicates of three independent experiments. IGR, intergenic region; GPC, glycoprotein complex; GFP, green fluorescent protein; LUC, luciferase; NP, nucleoprotein; ^****^*p*<0.001.

Pseudovirions of Ebola Zaire (pEBOV), Lake Victoria Marburg (pMARV), and Vesicular Stomatitis Virus (pVSV-G) containing a LUC reporter gene were produced in HEK-293T as described before (Cooper et al., 2020). The non-pathogenic replication-competent recombinant Cedar virus (rCedV) expressing LUC was propagated and titrated in Vero cells by PFU assay (Amaya et al., 2021b).

### 2.2. Cell-based antiviral assays

First, A549 cells infected with rPICV in 96- and 384-well plates were used to optimize the assay conditions for the antiviral assays. Then, A549, HeLa, BHK-21 and Vero cells were used for the HTS assays and/or to evaluate the antiviral activity of the hit compounds.

Cells were infected at MOIs 0.01-0.05 for assays with rPICV, rLCMV and rCedV, or at MOIs 0.5-0.8 for assays with LASV and JUNV. The replication of rPICV and rLCMV was quantified by measuring the LUC enzymatic activity in the infected cells 24 h post infection (hpi) or 48 hpi for rCedV. Briefly, the infected cells were lysed using the Neolite Reporter Gene Assay System (Revvity) and the LUC activity was examined after a 5-minutes incubation at room temperature by measuring the luminescent signal on an EnVision plate reader (PerkinElmer). Infection with LASV or JUNV was determined 36 hpi by immunostaining the infected cells with virus specific anti-nucleoprotein (NP) primary antibodies (Anantpadma et al., 2016). Stained cells were imaged using an Operetta high-content imaging system. Ribavirin at 10 µM was used as a positive control in all antiviral assays.

### 2.3. Automated high-throughput screenings (HTS) and cytotoxicity evaluation

The MicroSource Spectrum Collection, which is a chemically diverse small molecule library comprised of approximately 2,700 approved drugs, was used for the primary HTS. Compounds from this library were prepared in 384-well plates with the high-throughput liquid handler in a tissue-culture biosafety cabinet (PerkinElmer JANUS Automated Workstation). Compounds at a final concentration of 10 µM and rPICV were added to A549 cells also prepared in 384-well plates. After 24 h, the LUC activity was measured as described in subsection 2.2.

The potential hits selected from the primary HTS (as defined in the subsection 3.2) were re-screened for rPICV inhibition and tested for cytotoxicity in non-infected A549 cells at 10 µM. Viral replication was quantified through LUC activity evaluation and cell viability was determined using CellTiter-Glo (Promega) after 24 h of incubation.

### 2.4. Dose-response analyses of the prioritized hit compounds for antiviral activity determination and cytotoxicity evaluation

A dose-response analyses in different cell lines were employed for the prioritized hit compounds (as defined in the subsection 3.3) for the determination of the 50% cytotoxicity (CC_50_), 50% effective concentration (EC_50_), and the selectivity indexes (SI=CC_50_/EC_50_) against the arenaviruses, the pseudotyped viruses and rCedV. Infected cells were treated or not with 9-12 different concentrations of the compounds. Viral replication (or viral entry in the case of the pseudoviruses) and cell viability were assessed from 24-36 hpi as described in subsections 2.2 and 2.3.

### 2.5. Time-of-Addition assay (TOA)

BHK-21 cells infected with rPICV were treated with various concentrations of the top hit compounds at different time points as follows: 1h before the infection (−1 h), at the same time of infection (0 h), and at 2, 6 or 16 hpi (2 h, 6 h, and 16 h). rPICV replication was quantified by LUC activity measurement 24 hpi (subsection 2.3).

## 3. RESULTS

### 3.1. Assay optimization in 96- and 384-well plates for high-throughput screening (HTS)

A tri-segmented recombinant PICV (rPICV) expressing LUC was generated and used as a surrogate of highly pathogenic arenaviruses for HTS. The rPICV is a replication-competent virus (Dhanwani et al., 2016b) generated by splitting the S RNA genome segment into two segments (S1 and S2) and cloning the LUC and GFP reporter genes into the respective segments as shown in Fig. 1A.

rPICV’s ability to infect and replicate in A549 cells was firstly assessed, since the epithelium is the first barrier against infections and epithelial respiratory cells play an important role in arenavirus infection and propagation in the host (Brisse and Ly, 2019; Mpingabo et al., 2020). A549 cells in 96- and 384-well plates were infected with rPICV and viral replication was quantified by measuring the LUC activity in infected cells 24 hpi. Ribavirin, a known inhibitor of arenaviral RNA synthesis (Moreno et al., 2012), was used at 10 µM to evaluate whether it would serve as a positive control for the reduction of rPICV LUC activity.

In 96- and 384-well plate formats, infection with rPICV generated LUC luminescent signals higher than 3×10^5^ RLU (Fig. 1B and 1C), which led to high signal-to-background ratios (S/B >100) (Table 1). Ribavirin at 10 µM significantly reduced by 67% and 62% rPICV LUC activity in 96- and 384-well formats (Fig. 1B and 1C), respectively, indicating that the LUC expression in the infected cells is proportional to viral replication and that ribavirin can be used as the positive control in the assays. Both assay formats achieved robust Z’ and coefficients of variation (CV) values, Z’ ≥0.5 which corresponds to good assay sensitivity and low signal deviation within the assay (CV ≤20%) (Inglese et al., 2007; Zhang et al., 1999, Iversen et al., 2004) (Table 1), indicating that the assay conditions are suitable for HTS-based antiviral identification.

**Table 1.**
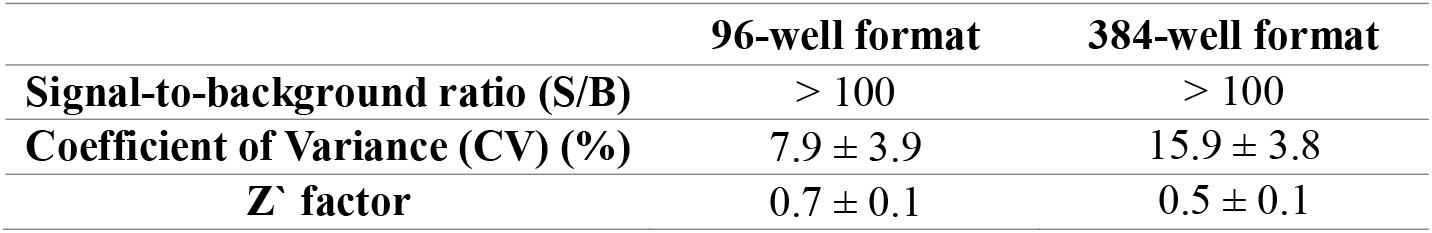
Parameters of rPICV-based HTS assay in 96- and 384-well formats.

### 3.2. HTS assays for the identification of rPICV small molecule inhibitors

A library of 2,700 small molecules plated in 384-well plates in a single concentration of 10 µM was screened for rPICV inhibition in A549 cells, as illustrated in Fig. 2A. The average Z’ score values for this primary screen was 0.5 ± 0.1, the average CV was 15.9% ± 3.8%, and S/B ratios for all plates were greater than 100. A 70% LUC activity reduction (or inhibition of rPICV replication) was used as the cutoff for considering compounds as potential hits (Fig. 2B). One hundred thirty-four compounds were cherry-picked as potential hit compounds, yielding a 4.3% hit rate (Fig. 2B).

**Fig. 2.**
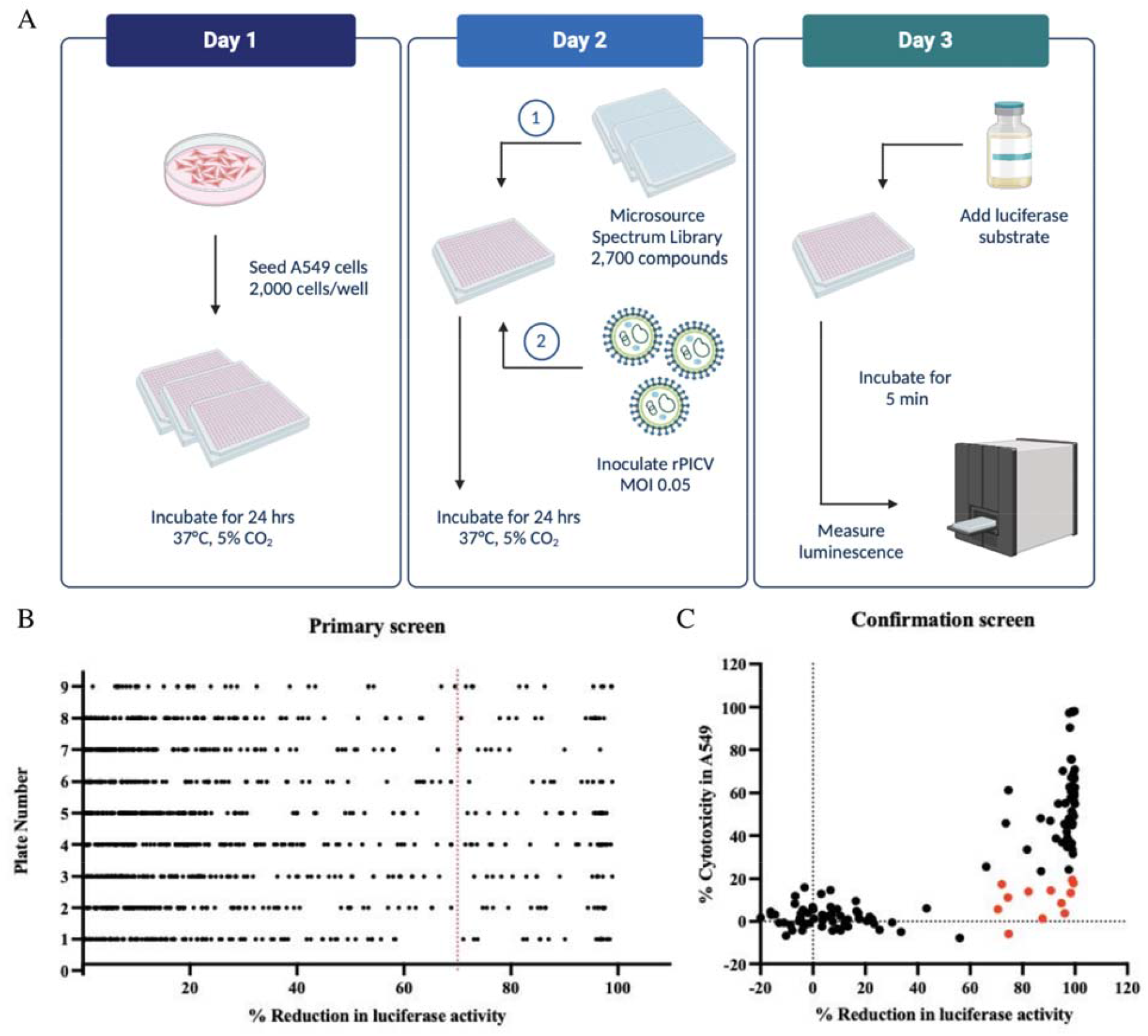
HTS workflow and criteria for hit selection. (A) HTS assay workflow in 384-well format. At day 1, A549 cells were seeded into 384-well plates at a density of 2,000 cells/well and incubated at 37°C and 5% CO_2_ for 24 h. The day after (day 2), the small molecule library also prepared in 384-well plates was added to A549 cell plates at a final concentration of 10 µM. Cells were then infected with rPICV at an MOI of 0.05. After a further 24 h incubation, at day 3, Neolite Reporter Gene Assay System was added and LUC activity was measured by luminescence reading. (B) Overview of the primary HTS results of the 2,700-small molecule library. Each black dot represents one compound along with its percentage of reduction in LUC activity, shown in the X axis. One hundred thirty-four compounds exhibited ≥70% reduction in LUC activity (or ≥70% reduction of rPICV replication) and were considered potential hits. (C) Confirmation screen results. The 134 potential hits (presented as dots) were retested at 10 µM for anti-rPICV activity and for cytotoxicity to non-infected A549 cells. Twelve compounds highlighted in red displayed reduction in rPICV replication over 70% and cytotoxicity to A549 cells lower than 20%.

For the confirmation screen, the 134 potential hits were retested at 10 µM for anti-rPICV activity. A single dose (10 µM) cytotoxicity assay was performed in parallel to rule out false positive inhibition results due to compound’s cytotoxicity. The antiviral screening plate presented a Z’ score of 0.5 and a CV of 15.1%. The Z’ and CV values for the cytotoxicity assay were 0.9 and 3.8%, respectively. Both plates showed S/B ratios >1000. As a result of this screen, twelve compounds qualified as “confirmed hits”, since they showed >70% inhibition of rPICV replication and <20% cytotoxicity at 10 µM (Fig. 2C). Interestingly, ribavirin met these criteria as a hit, showing ∼75% inhibition of viral replication and no cytotoxicity to A549 cells at 10 µM (Table S1). This result highlights the robustness of the HTS assays. Besides ribavirin, eight other hit compounds were prioritized for further evaluation based on the lack of potential toxicophores (Table S1) (Daina et al., 2017). Three of the confirmed hits were deprecated based on the presence of potential toxicophores and other poor drug-like properties, and were not explored further (Table S2).

### 3.3. Validation of the antiviral activity of the prioritized hits against arenaviruses

To validate the antiviral activity of the eight prioritized hit compounds listed in Table S1 and to evaluate their potential as broad-spectrum anti-arenaviruses inhibitors, they were tested against representatives of NW (rPICV and LASV (Josiah)) and OW arenaviruses (rLCMV and JUNV (Romero)). Dose-response curves were performed to determine the potency and selectivity of those hits (including ribavirin as a positive control) for rPICV inhibition in A549 cells. The four top hits highlighted in bold in Table 2, which are 2’,4’-dihydroxychalcone, gossypol, lapachol, and piplartine, inhibited rPICV replication with EC_50_ values in the low micromolar range (<5 µM) and with better potency than ribavirin (EC_50_=6.6 µM). These top hit compounds showed mid to low cytotoxicity to A549 cells (CC_50_ values ranging from 36 to >100 µM), and good SIs (>10) (Table 2). Despite the low cytotoxicity found for bronopol, DIDS sodium salt, phenothiazine and podofilox (CC_50_ ≥100 µM), these molecules showed poor or no inhibition of viral replication, with EC_50_ values >30 µM and, consequently, low SIs (Table 2).

**Table 2.** Pharmacological parameters of the eight prioritized hit compounds and ribavirin for rPICV and rLCMV in different cell lines.

| Compounds | A549 |  |  | HeLa |  |  | BHK-21 |  |  |  |  |
| --- | --- | --- | --- | --- | --- | --- | --- | --- | --- | --- | --- |
|  | CC<sub>50</sub> | rPICV<br>EC<sub>50</sub> | rPICV<br>SI | CC<sub>50</sub> | rPICV<br>EC<sub>50</sub> | rPICV<br>SI | CC<sub>50</sub> | rPICV<br>EC<sub>50</sub> | rPICV<br>SI | rLCMV<br>EC<sub>50</sub> | rLCMV<br>SI |
| **2',4'-**<br>**Dihydroxychalcone** | **> 100** | **1.2 ± 0.2** | **83** | **41.8 ± 1.0** | **3.9 ± 0.1** | **11** | **> 100** | **5.6 ± 0.4** | **> 18** | **10.6 ± 1.1** | **> 9** |
| **Gossypol** | **61.8 ± 3.7** | **1.6 ± 0.6** | **39** | **> 100** | **6.1 ± 0.1** | **> 16** | **> 100** | **2.2 ± 0.06** | **> 45** | **7.3 ± 0.8** | **> 14** |
| **Lapachol** | **> 100** | **3.8 ± 0.8** | **> 26** | **77.7 ± 3.1** | **3.6 ± 0.08** | **22** | **> 100** | **3.6 ± 0.08** | **> 28** | **3.7 ± 0.2** | **> 27** |
| **Piplartine** | **35.9 ± 2.6** | **2.5 ± 0.6** | **14** | **> 100** | **5.4 ± 0.1** | **> 18** | **> 100** | **4.7 ± 0.02** | **> 21** | **6.7 ± 0.1** | **> 15** |
| Ribavirin | > 100 | 6.6 ± 0.6 | > 15 | > 100 | 2.4 ± 0.2 | > 42 | > 100 | 2.6 ± 0.2 | > 38 | 3.4 ± 0.01 | > 29 |
| Bronopol | > 100 | 31.0 ± 1.8 | > 3 | ND | ND | ND | ND | ND | ND | ND | ND |
| DIDS Sodium Salt | > 100 | > 100 | unk | ND | ND | ND | ND | ND | ND | ND | ND |
| Phenothiazine | ~ 100 | 34.0 ± 10.8 | 3 | ND | ND | ND | ND | ND | ND | ND | ND |
| Podofilox | > 100 | NI | unk | ND | ND | ND | ND | ND | ND | ND | ND |

EC<sub>50</sub> and CC<sub>50</sub> values are presented as mean ± SD in μM; SI, selectivity index (CC<sub>50</sub>/EC<sub>50</sub>); unk, unknown; NI, no inhibition; ND, not determined.

The anti-rPICV activity of the four top hits (2’,4’-dihydroxychalcone, gossypol, lapachol, and piplartine) was validated in two more cell lines, HeLa and BHK-21. The compounds also showed mild to low cytotoxicity in HeLa and BHK-21 (CC_50_ values from 42 to >100 µM), and inhibited rPICV replication in both cell lines with low EC_50_ values (≤6 µM) and good SIs (>10) (Table 2). The inhibitory activity of these compounds was further confirmed using another LUC-expressing recombinant prototypic arenavirus, rLCMV. The potency of the compounds in inhibiting rLCMV replication in BHK-21 cells was slightly lower than that of rPICV, but the EC_50_ values were still within the micromolar range (≤10 µM) and the SIs >10 (Table 2).

As a proof-of-concept, two of the top hit compounds showing consistent inhibitory results against rPICV and rLCMV and more favorable SI values across the different cell lines, gossypol and lapachol (Table 2), were also tested against LASV (Josiah) and JUNV (Romero) in A549 cells. The Z’ score for these assays varied from 0.63 to 0.96 and the average signal-to-noise ratio (S/N) was >100. After a 36-h incubation period, lapachol’s cytotoxicity was low (CC_50_ >50 µM) and its EC_50_ values for LASV and JUNV were of 10 and 14 µM, respectively, which led to reasonable SIs (Fig. 3 and Table 3). Gossypol showed increased cytotoxicity after 36 h, when compared with 24 h of treatment (CC_50_=28 µM, 2 times higher than 24 h time point) (Fig. 3 and Table 3). Therefore, despite high potencies of inhibition of LASV and JUNV replication in the low micromolar range (EC_50_ ≤5 µM), gossypol’s selectivity was lower than 10 (Fig. 3 and Table 3). Ribavirin was used as the positive control and inhibited LASV and JUNV replication with good selectivity, showing SI values >10 (Table 3).

**Table 3.** Pharmacological parameters of the top hit compounds for the laboratory strains of LASV (Josiah) and JUNV (Romero) in A549 cells.

| **Compounds** | **CC<sub>50</sub>** | **LASV**<br>**EC<sub>50</sub>** | **LASV**<br>**SI** | **JUNV**<br>**EC<sub>50</sub>** | **JUNV**<br>**SI** |
| --- | --- | --- | --- | --- | --- |
| Gossypol | 28.4 ± 5.8 | 4.6 ± 0.4 | 6 | 5.2 ± 0.2 | 5 |
| Lapachol | > 50 | 9.7 ± 0.6 | > 5 | 14.0 ± 0.3 | > 4 |
| Ribavirin | > 500 | 25.3 ± 1.1 | > 20 | 44.4 ± 3.3 | > 11 |

EC<sub>50</sub> and CC<sub>50</sub> values are presented as mean ± SE in μM; SI, selectivity index (CC<sub>50</sub>/EC<sub>50</sub>).

**Fig. 3.**
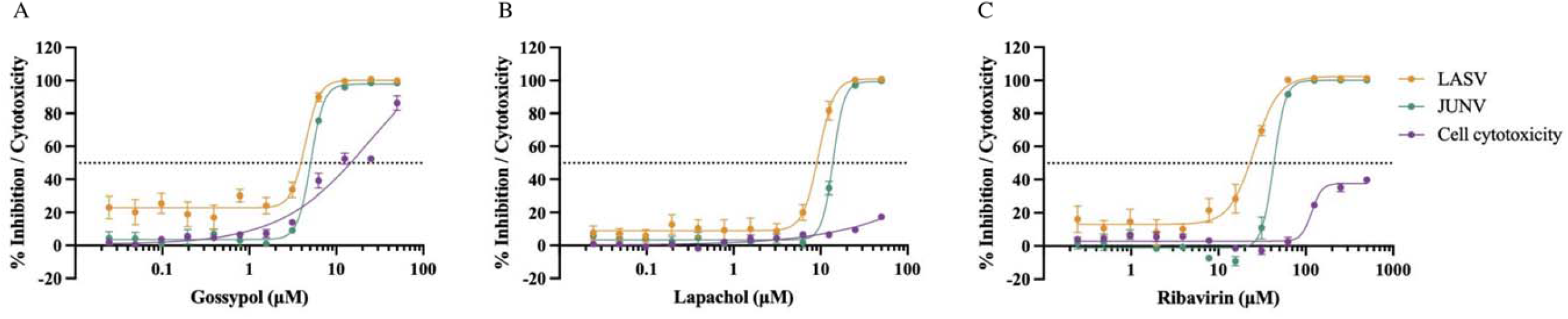
Antiviral activity and cytotoxicity of two top hit compounds and ribavirin against laboratory strains of LASV (Josiah) and JUNV (Romero). A549 cells in 384-well plates were pretreated with various concentrations of gossypol (A), lapachol (B) or ribavirin (C) for 1h at 37°C, then infected with LASV or JUNV (MOI of 0.5-0.8) under BSL-4 containment conditions. After 36 h, cells were fixed with 10 % formalin solution and incubated with anti-LASV or anti-JUNV nucleoprotein (NP) specific primary antibodies followed by another incubation with fluorophore conjugated secondary antibody and Hoechst 33342 for nuclear staining. Images of infected cells were acquired on an Operetta high-content imaging system followed by analysis using Harmony software. Data were normalized by the untreated controls for the calculation of the percentage of inhibition/cytotoxicity, which were then used to obtain the pharmacological parameters of CC_50_, EC_50_ and SIs. The purple dose-response curves represent the percentage of cytotoxicity of each compound to A549 cells. The orange and green dose-response curves represent the percentage of inhibition of LASV and JUNV replication, respectively, of each compound. The data represent as mean ± SE from at least three replicates of two independent experiments.

The antiviral activity data presented in Tables 2 and 3 are important to validate the HTS data using rPICV as a surrogate for pathogenic arenaviruses. The selection of the direct-acting antiviral ribavirin as well as the identification of host-directed drugs (e.g. lapachol, Table S1) that inhibited different strains of arenaviruses demonstrate the feasibility of this HTS strategy to uncover broad-spectrum small molecule inhibitors against arenaviruses.

### 3.4. Insights into the mechanism of action of the top hit compounds

The four top hits, 2’,4’-dihydroxychalcone, gossypol, lapachol, and piplartine, have been previously shown to exhibit various biological activities including host-directed antiviral activity through the modulation of cellular pathways important for viral replication and virus-directed antiviral activity through the inhibition of viral proteins such as viral polymerases (Table S1). To evaluate the specificity of inhibition by the top hits towards arenaviruses and to investigate whether they could be targeting cellular pathways, compounds were tested against viruses belonging to families other than *Arenaviridae*, such as filoviruses (pEBOV and pMARV), rhabdovirus (pVSV-G), and henipavirus (rCedV).

As expected, ribavirin, a broad-spectrum inhibitor of viral RNA synthesis used as a positive control, did not impair the entry/fusion of any pseudovirus in A549 cells (Table 4). On the other hand, this compound reduced rCedV replication in Vero76 cells with an EC_50_ of ∼17 µM (Table 4). Interestingly, 2’,4’-dihydroxychalcone, gossypol, and lapachol inhibited pEBOV, pMARV, and pVSV-G infection, with similar EC_50_ values across the pseudoviruses and good SIs (Table 4). The 2’,4’-dihydroxychalcone and lapachol compounds also showed reasonable antiviral activity against rCedV (Table 4). These data suggest that these three compounds could be targeting either early stages of viral replication or early antiviral response triggered in the host cells. Piplartine was considerably more cytotoxic to A549 and to Vero76 cells after 48 h of treatment, with CC_50_ of 12-13 µM, and showed weak inhibition of the tested viruses (EC_50_ of 6-28 µM, Table 4), leading to lower SI values than those obtained for rPICV and rLCMV (Table 2). These data suggest specific inhibition of arenaviruses replication.

**Table 4.** Antiviral profile of the top hit compounds against pEBOV, pMARV, pVSV and rCedV.

| Compounds | A549 |  |  |  |  |  |  | Vero 76 |  |  |
| --- | --- | --- | --- | --- | --- | --- | --- | --- | --- | --- |
|  | CC<sub>50</sub> | pEBOV<br>EC<sub>50</sub> | pEBOV<br>SI | pMARV<br>EC<sub>50</sub> | pMARV<br>SI | pVSV<br>EC<sub>50</sub> | pVSV<br>SI | CC<sub>50</sub> | rCedV<br>EC<sub>50</sub> | rCedV<br>SI |
| 2',4'-<br>Dihydroxychalcone | > 100 | 3.8 ± 0.2 | > 26 | 3.9 ± 0.2 | > 26 | 3.8 ± 0.3 | > 26 | > 100 | 20.9 ± 3.0 | > 5 |
| Gossypol | 14.8 ± 0.8 | 1.2 ± 0.3 | 12 | 1.7 ± 0.2 | 9 | 0.6 ± 0.07 | 25 | 15.2 ± 1.2 | 4.7 ± 0.3 | 3 |
| Lapachol | 48.8 ± 1.0 | 1.2 ± 0.5 | 40 | 1.2 ± 0.2 | 40 | 3.6 ± 0.4 | 13 | > 100 | 11.1 ± 0.7 | > 9 |
| Piplartine | 13.2 ± 0.4 | 26.4 ± 2.8 | 0.5 | 27.9 ± 0.2 | 0.5 | 14.1 ± 0.4 | 1 | 11.6 ± 1.3 | 6.2 ± 0.9 | 2 |
| Ribavirin | >100 | NI | unk | NI | unk | NI | unk | > 100 | 16.8 ± 1.3 | > 6 |

EC<sub>50</sub> and CC<sub>50</sub> values are presented as mean ± SD in µM; SI, selectivity index (CC<sub>50</sub>/EC<sub>50</sub>); NI, no inhibition; unk, unknown.

To better understand the potential temporal pattern of inhibition mediated by the top hit compounds, time-of-addition assays using rPICV were performed. Briefly, serially diluted compounds were added to infected BHK-21 cells at different time points, 1 h before the infection (−1 h), at the time of infection (0 h), or at 2, 6, or 16 hpi, which is the approximate timing for one round of rPICV replication. Viral replication was measured 24 hpi, and the potency of inhibition of the compounds was calculated for each time point. In accordance with its mechanism of inhibition of viral RNA replication, the treatment with ribavirin could be efficiently delayed up to 6 h after infection without loss of activity, as evidenced by the 3-fold increase in its EC_50_ value when added 16 hpi (Table 5). The same inhibition profile was observed for 2’,4’-Dihydroxychalcone, lapachol and piplartine, suggesting that these compounds target steps at a similar stage as the viral RNA synthesis (Table 5). Gossypol’s antiviral activity was maintained up to the time of 2 h of drug addition, suggesting its effect during earlier stages of the viral replication cycle (Table 5).

**Table 5.**
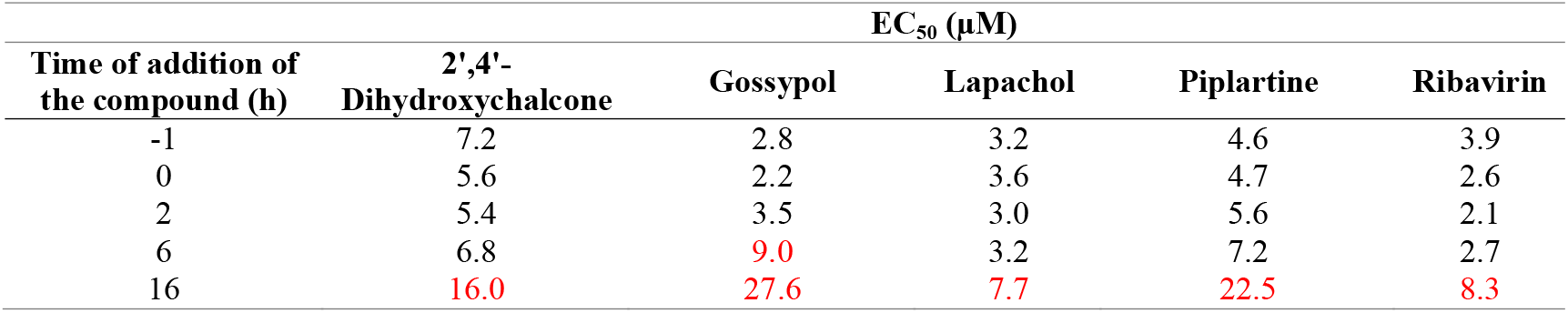
Potency of inhibition of the top hit compounds in time-of-addition assay.

| Time of addition of<br>the compound (h) | EC<sub>50</sub> (μM) |  |  |  |  |
| --- | --- | --- | --- | --- | --- |
|  | 2',4'-<br>Dihydroxychalcone | Gossypol | Lapachol | Piplartine | Ribavirin |
| -1 | 7.2 | 2.8 | 3.2 | 4.6 | 3.9 |
| 0 | 5.6 | 2.2 | 3.6 | 4.7 | 2.6 |
| 2 | 5.4 | 3.5 | 3.0 | 5.6 | 2.1 |
| 6 | 6.8 | 9.0 | 3.2 | 7.2 | 2.7 |
| 16 | 16.0 | 27.6 | 7.7 | 22.5 | 8.3 |

## 4. DISCUSSION

In this work, the replication-competent rPICV, a non-pathogenic NW arenavirus, was successfully utilized for the screening and identification of broad-spectrum small molecule inhibitors against arenaviruses under a BSL-2 laboratory condition. The use of rPICV expressing LUC as a reporter gene was adapted to 384-well assay format with Z’ ≥0.5, CV ≤20%, and high S/B, demonstrating the feasibility of the HTS-based antiviral identification through a sensitive and robust assay.

Since it was the first time that rPICV was being used as a tool for HTS, a cutoff of 70% reduction in the LUC expression in infected cells was set for the primary screen and 134 potential hits were identified from a library of 2,700 approved small molecules, providing a high hit rate of 4.3%; usually, the expected hit rate for HTSs is ∼1% (Dreiman et al., 2020). As a way to narrow down the number of active compounds and to avoid the selection of false positive hits due to cell toxicity, the anti-rPICV activity of the potential hits was reconfirmed, along with an evaluation of their cytotoxicity. Five top hits, which included ribavirin (a nucleoside analog with known anti-arenaviral activity), were selected from this confirmation screen and indeed showed good potency (EC_50_ <5 µM) and high selectivity (SI >10) in inhibiting the replication of rPICV in human and non-human cells lines, and of rLCMV (an OW prototypic arenavirus) with comparable parameters. The antiviral activity of two top hits was also validated against other OW and NW arenaviruses, LASV and JUNV, respectively. These data demonstrated the efficacy of using the rPICV-based HTS in BSL-2 to identify relevant antiviral compounds against BSL-3/4 arenavirus pathogens.

Drug discovery platforms for highly pathogenic BSL-3/4 arenaviruses suitable for BSL-2 facilities mostly rely on strategies that target specific steps of the virus life cycle. For instance, pseudotyped arenaviruses expressing the envelope glycoprotein have been used for the identification of entry/fusion inhibitors (Iyer et al., 2024), while minireplicon systems were developed for the evaluation of molecules inhibiting viral RNA transcription and replication (Droniou-Bonzom et al., 2011; Mendenhall et al., 2010), and virus-like particles produced by self-budding of arenaviruses matrix protein (Z) for the study of budding inhibitors (Lu et al., 2014). Otherwise, the use of authentic recombinant arenaviruses in cell-based assays, such as rLCMV (Ngo et al., 2015) and rPICV (described in this study), which can replicate in different cell lines, allow the identification of inhibitors targeting several steps during viral life cycle and/or host-cell pathways involved in viral multiplication. Among the five top hits identified in this work, 2’,4’-dihydroxychalcone, lapachol, piplartine, and ribavirin appeared to act at a later stage of the viral life cycle than gossypol (Table 5). Specifically, ribavirin, known to inhibit arenaviral RNA synthesis (Moreno et al., 2012). Thus, the rPICV-based HTS developed in this study demonstrated its efficacy in identifying antiviral compounds with diverse mechanisms of action. Although further mechanistic studies are necessary for target determination, it appears that the hits identified using rPICV might have host- and virus-directed antiviral activities.

In summary, the rPICV-based HTS assay described herein has proven to be a suitable and safe surrogate for the identification of broad-spectrum small molecule inhibitors against arenaviruses with a variety of antiviral mechanisms. Future studies may include screening of larger libraries, which may lead to the identification of compounds with good drug-like characteristics that can be chemically derivatized and optimized for further development as potential anti-arenaviral therapeutics.

## Supporting information

Supplemental Methodology

## ACKNOWLEDGMENTS

We thank K.K. Conzelmann (Ludwig-Maximilians-Universität, Germany) for the BSRT7-5 cells and Juan de la Torre (the Scripps Research Institute) for the LCMV reverse genetics system. We also acknowledge Michael Flint, Christina Spiropoulou, and Jamie Kelly (Viral Special Pathogens Branch, Centers for Disease Control and Prevention, CDC) for BSL-4 assays.

## AUTHOR CONTRIBUTIONS

**Carolina Q. Sacramento:** Methodology, Investigation, Formal Analysis, Writing – original draft, Writing – review and editing; **Ryan Bott:** Methodology, Investigation, Formal Analysis, Writing – review and editing; **Qinfeng Huang:** Methodology, Investigation, Formal Analysis; **Brett Eaton:** Methodology, Investigation, Formal Analysis; **Elena Postnikova:** Methodology, Investigation, Formal Analysis, Writing – review and editing; **Ahmad J. Sabir:** Methodology, Investigation, Formal Analysis; **Malaika D. Argade:** Methodology, Investigation, Formal Analysis, Writing – review and editing; **Kiira Ratia:** Investigation; **Manu Anantpadma:** Formal Analysis, Supervision, Writing – review and editing; **Paul R Carlier:** Project Administration, Resources, Supervision, Writing – review and editing; **Hinh Ly:** Conceptualization, Project Administration, Resources, Supervision, Writing – review and editing; **Yuying Liang:** Conceptualization, Project Administration, Resources, Supervision, Writing – review and editing; **Lijun Rong:** Conceptualization, Project Administration, Resources, Supervision, Writing – review and editing.

## FUNDING

All the work done at the IRF-Frederick was funded in whole or in part with federal funds from the National Institute of Allergy and Infectious Diseases, National Institutes of Health, Department of Health and Human Services, under Contract No. HHSN272201800013C. B.E., M.A. performed this work as employees of Laulima Government Solutions, LLC. Subcontractors to Laulima Government Solutions, LLC who performed this work are: E.P. an employee of Tunnell Government Services, Inc.

## DISCLAIMER

The content of this publication does not necessarily reflect the views or policies of the US Department of Health and Human Services (DHHS) or of the institutions and companies affiliated with the authors.

## DECLARATION OF INTERESTS

The authors have declared no conflict of interest.

## Notes

### Competing Interest Statement

The authors have declared no competing interest.

### Summary of Updates

A following disclaimer was added to the manuscript: "The content of this publication does not necessarily reflect the views or policies of the US Department of Health and Human Services (DHHS) or of the institutions and companies affiliated with the authors".

