## Supplemental Methodology for "Recombinant Pichinde reporter virus as a safe and suitable surrogate for high-throughput antiviral screening against highly pathogenic arenaviruses"

### Supplementary Materials and Methods

#### 1.1. Cells and viruses

A549, HEK-293T, HeLa, BHK-21 and Vero cell lines were maintained in high glucose Dulbecco's Modified Eagle Media (DMEM) medium supplemented with 100 µg/mL streptomycin, 100 U penicillin (P/S) and 10% fetal bovine serum (FBS) at 37°C and 5% $CO_2$. BSRT7-5 cells are BHK-21 cells stably expressing T7 RNA polymerase, and were obtained from K.K. Conzelmann (Ludwig-Maximilians-Universität, Germany) and grown in minimal essential medium (MEM) supplemented with 10% FBS, 1 µg/ml Geneticin, and 50 µg/ml P/S.

Tri-segmented recombinant PICV (rPICV) expressing firefly luciferase (LUC) and green fluorescent protein (GFP) was generated by splitting the S RNA genome segment into two (S1 and S2), as previously described (Dhanwani et al., 2017). The LUC and GFP genes were cloned into new multiple cloning sites in the S2 and S1 genomic segments, respectively (Fig. 1A). The S1-, S2- and L-encoding plasmids were transfected into BSRT7-5 cells. Infectious rPICV expressing LUC and GFP was collected from the supernatants, propagated in BHK-21 cells and titrated in Vero cells by plaque forming unit (PFU) assays (Dhanwani et al., 2017, 2016). rLCMV expressing LUC and GFP was created in a similar approach, using plasmids from the LCMV reverse genetics system provided by The Scripps Research Institute. Laboratory manipulation of rPICV and rLCMV under BSL-2 conditions was approved by the Institutional Biosafety Committees at the University of Minnesota and University of Illinois Chicago in accordance with NIH guidelines.

All work with the laboratory strains of LASV (Josiah) and JUNV (Romero) was conducted in a BSL-4 laboratory at the Integrated Research Facility Frederick (IRF Frederick), in compliance with local, state, and federal regulations for handling select agents. Working stocks of LASV

(Josiah) and JUNV (Romero) were generated by infecting Vero cells at a multiplicity of infection (MOI) of 0.01-0.1, followed by incubation for 5 to 7 days. The viruses from the supernatants were clarified by centrifugation at 2,500x*g*, and then aliquoted into single-use vials for storage at -70°C until use.

**1.2. Compound library**

The MicroSource Spectrum Collection is a chemically diverse small molecule library comprised of approximately 2,700 FDA and/or internationally approved drugs (Asia and Europe) and was provided by the University of Illinois Chicago Research Resources Center (UIC RRC). Ribavirin and the prioritized hits selected from the screenings were purchased from MedChemExpress. Compound identity and purity were confirmed in each case by $^{1}$H nuclear magnetic resonance spectroscopy and high-performance liquid chromatography-mass spectrometry. The master stocks of all compounds were prepared in DMSO and further diluted in culture media in a way that the DMSO concentration never exceeds 1%.

**1.3. Optimization of the cell-based antiviral assays**

A549 cells in white opaque 96-well plates (5,000 cells/well) were infected with rPICV at the MOI of 0.01 for 1h at 37ºC. After incubation, the inoculum was removed and the cells were treated with 10 µM of ribavirin or untreated. For the assay in 384-well format, A549 cells (2,000 cells/well) were treated with ribavirin (10 µM) right before the infection with rPICV at the MOI of 0.05. Cells were further incubated for 24 h and lysed using the Neolite Reporter Gene Assay System (Revvity), according to manufacturer's instructions. The LUC enzymatic activity was measured after a 5 minutes-incubation at room temperature by reading the luminescent signal on an EnVision plate reader (PerkinElmer). In both assay formats, wells containing cells only (mock) and cells with virus (negative controls) were used for the determination of quality parameters such as Z' score, signal-to-background ratio (S/B), and coefficient of variation (CV).

#### 1.4. Automated primary high-throughput screening (HTS)

Compounds from the testing library prepared in 384-well plates were diluted in screening medium (phenol red-free DMEM with 2% FBS and 2% L-glutamine) with the high-throughput liquid handler in a tissue-culture biosafety cabinet (PerkinElmer JANUS Automated Workstation) and added in a final concentration of 10 μM to A549 cells. rPICV (MOI 0.05) was then added to the cell plates. Ribavirin at 10 μM was used as the positive control. After 24 h, the LUC activity was measured as described in subsection 1.3. 70% reduction of the LUC activity (which corresponds to a 70% reduction of rPICV replication) was used as the criterion for designating a compound as a "potential hit". Mock and negative control wells were included in each plate for the calculation of assay quality parameters on a plate basis.

#### 1.5. Confirmation antiviral screening and cytotoxicity evaluation

The potential hits were re-screened for rPICV replication inhibition and tested for cytotoxicity in non-infected A549 cells at 10 μM. A549 cells in 384-well plates were treated with 10 μM of the potential hits followed by rPICV inoculation (MOI 0.05) or the addition of screening medium for the cytotoxicity assay. After 24h, Neolite or CellTiter-Glo (Promega, used for the determination of cell viability) was added to the corresponding plates. The luminescent signal was measured as described in the subsection 1.3. Confirmed hits were defined as those compounds showing rPICV replication inhibition over 70%, and cell cytotoxicity below 20%.

#### 1.6. Dose-response analysis of prioritized hit compounds using recombinant arenaviruses

A 9-point dose-response analysis was employed for the prioritized hit compounds for the determination of the half-maximum effective activity ($EC_{50}$), the 50% cytotoxicity concentration ($CC_{50}$) and the selectivity indexes ($SI=CC_{50}/EC_{50}$) against rPICV and rLCMV. A549, HeLa or BHK-21 cells in 96-well plates were infected or not infected with rPICV or rLCMV (MOIs 0.01 to 0.05) for 1h at 37°C. The inoculum were removed from the infected plates and the cells were treated with different concentrations of the compounds. Culture medium with 1% DMSO was used as a negative control. After 24h, Neolite or CellTiter-Glo was added to the plates for luminescence measurement (subsection 1.3). The data was normalized to the negative control to acquire percentage inhibition or cytotoxicity values. Normalized values were plotted to determine of $CC_{50}$ and $EC_{50}$ values.

#### 1.7. Inhibition assays with laboratory strains of infectious LASV and JUNV

A549 cells in 384-well plates were pretreated with 12 different concentrations of two of the top hit compounds for 1 h at 37°C. Plates were then infected with LASV (Josiah) or JUNV (Romero) at MOIs from 0.5 to 0.8. After 36 h, the cells were fixed with a 10% formalin solution. Infection was determined through techniques previously described (Anantpadma et al., 2016). Briefly, the cells were stained with virus-specific anti-nucleoprotein (NP) primary antibodies (anti-LASV NP, Cambridge Bio, 1:500 dilution or custom-made anti-JUNV NP, 1:1000), followed by staining with fluorophore-conjugated secondary antibodies: Goat anti-Mouse IgG Alexa Fluor™ 488 for LASV, or Goat anti-Rabbit IgG Alexa Fluor™ 647 for JUNV (both from ThermoFisher Scientific, 1:2500). Hoechst 33342 (1:2500 dilution) was used for nuclear staining. Stained cells were imaged using an Operetta high-content imaging system and analyzed with Harmony software. One method to estimate cell viability and determine the compounds' cytotoxicity was by comparing the nuclei count from treated to uninfected controls. At the time of fixation of infected cells, a second plate was used to assess cytotoxicity using the CellTiter-Glo assay.

#### 1.8. Counter-screening of the hits using pseudotyped filoviruses, pseudotyped VSV and recombinant Cedar virus (rCedV)

The top hits were tested against pseudotyped Ebola Zaire (pEBOV), Lake Victoria Marburg (pMARV) (Cooper et al., 2020) and Vesicular Stomatitis Virus (pVSV), and LUC-expressing recombinant Cedar virus (rCedV) (Amaya et al., 2021). The pseudoviruses containing a LUC reporter gene were produced by transfecting HEK-293T cells as previously described (Cooper et al., 2020) with the following plasmids: EBOV-GP, MARV-GP or VSV-G and the HIV-1 pro-viral vector pNL4–3.Luc.R−E− (BEI Resources). A549 cells in 96-well plates were transduced with pseudoviruses and compounds at various concentrations. Pseudovirions with 1% DMSO were used as a negative control. The assay with rCedV was performed in Vero76 cells prepared in 96-well plates, infected with rCedV (MOI 0.01) for 1h at 37°C, and treated with different concentrations of the top hits after inoculum removal. After 48h of incubation, the LUC activity was measured as described in subsection 1.3.

#### 1.9. Time-of-addition assay (TOA)

BHK-21 cells in 96-well plates were infected with rPICV (MOI 0.05) and treated with different concentrations of the top hit compounds at different time points as follows: 1h before the infection (-1 h time point), at the same time of infection (0 h), and at 2, 6 or 16 h post-infection (hpi) (2 h, 6 h, and 16 h). rPICV replication was quantified by LUC activity measurement 24 hpi. Data were normalized by the untreated controls and the potency of inhibition ($EC_{50}$ values) of the compounds was calculated for each time point.

#### 2.10. Data analysis

Data are presented as the arithmetic mean and standard deviation (SD) or standard error (SE) of the values obtained from at least three experimental replicates, unless otherwise mentioned. Statistical analyses to calculate $EC_{50}$ and $CC_{50}$ values were performed with nonlinear regression four-parameters variable slope curves using GraphPad Prism software. Selectivity index (SI) was calculated as the ratio between $CC_{50}$ and $EC_{50}$ values.

The quality of the assays was evaluated through the calculation of $Z'$ scores, signal-to-background (S/B) ratios, signal-to-noise (S/N) ratios and coefficients of variance (CV) for each plate according to Zhang et al., 1999, considering the mock wells (cells only) and negative controls (rPICV-infected and untreated wells).
