## Supplemental Tables for "Recombinant Pichinde reporter virus as a safe and suitable surrogate for high-throughput antiviral screening against highly pathogenic arenaviruses"

### SUPPLEMENTARY TABLES

**Table S1.** List of prioritized hits with their chemical structure, previously reported biological activity and percentages of rPICV replication inhibition and of cytotoxicity to A549 cells in the primary and confirmation screenings.

| Compound | Chemical Structure | Biological Activity | Primary Screen Inhibition [a] | Confirmation Screen | |
|---|---|---|---|---|---|
| | | | | Inhibition [a] | Cytotoxicity [a] |
| **2',4'-Dihydroxychalcone** | 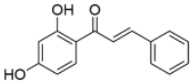 | Hsp90 inhibitor (Seo et al., 2015); host-directed effects and direct antiviral activity against SARS-CoV-2 and MERS, influenza viruses, HIV-1, HSV-1/2, HBV/HCV, DENV-2 and HCMV (Nematollahi et al., 2023) | 88% | 98% | 13% |
| **Bronopol** | 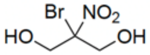 | Wide-spectrum antimicrobial properties (microbiocide/microbiostat) (Legin, 1996) | 85% | 95% | 8.5% |
| **DIDS sodium salt [b]** | 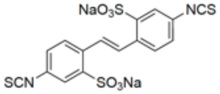 | Anion transport inhibitor (Santos et al., 1999); intracellular pH regulator (Lane et al., 1999) | 86% | 96% | 3.8% |
| **Gossypol** | 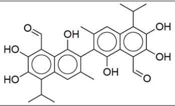 | Dehydrogenase inhibitor ; anticancer (Paunovic et al., 2023); direct antiviral activity against HIV-1 (Lin et al., 1989), HSV-2 (Bai et al., 2023), coronaviruses (Wang et al., 2022), ZIKV and DENV (Gao et al., 2019) | 97% | 74% | 11% |
| **Lapachol** | 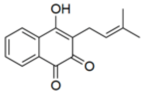 | Anticancer (Marques et al., 2020), antileshimanial, anti-inflammatory and bactericidal properties (Hussain et al., 2007); direct antiviral activity against HSV-1/2, HIV-1 and influenza (Hussain et al., 2007) | 90% | 99% | 20% |
| **Piplartine** | 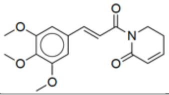 | Anticancer (Duarte et al., 2023); host-directed anti-SARS-CoV-2 (Tang et al., 2023) and anti-ZIKV effects (Lu et al., 2021) | 84% | 91% | 15% |
| **Phenothiazine** | 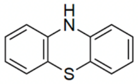 | Antipsychotic (modulates neurotransmitter activities), antibacterial and antineoplastic activities (Varga et al., 2017) | 80% | 88% | 1.4% |
| **Podofilox** | 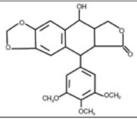 | Antineoplastic activity (Han et al., 2023); host-directed antiviral activity (Cohen et al., 2016)12/19/24 11:01:00 AM | 72% | 72% | 17% |
| **Ribavirin** | 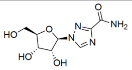 | Broad-spectrum direct antiviral activity through RNA synthesis inhibition (viral DNA/RNA polymerase inhibitor) (Nyström et al., 2019) | 73% | 75% | -5.9% |

[a] % inhibition and % cytotoxicity at 10 μM; [b] 4,4'-Diisothiocyanostilbene-2,2'-Sulfonic Acid Sodium Salt.

**Table S2.** List of deprecated hits with their chemical structures, reason(s) for disqualification, and percentages of rPICV replication inhibition and of cytotoxicity to A549 cells in the primary and confirmation screenings.

| Compound | Chemical Structure | Disqualifications | Primary Screen Inhibition [a] | Confirmation Screen | |
|---|---|---|---|---|---|
| | | | | Inhibition [a] | Cytotoxicity [a] |
| **Cedrelone** | 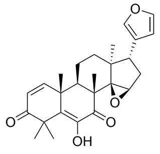 cedrelone | Potential toxicophore (epoxide) | 98% | 82% | 12% |
| **Gedunin** | 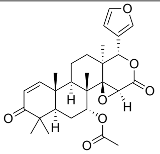 | Potential toxicophore (epoxide), poor aqueous solubility | 72% | 99% | 16% |
| **Erysolin** | 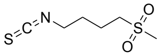 | Potential toxicophore (isothiocyanate) | 87% | 71% | 4% |

[a] % inhibition and % cytotoxicity at 10 μM.
